# Pressure protein denaturation compared to thermal and chemical unfolding - Analyses with cooperative models

**DOI:** 10.1101/2024.08.14.608019

**Authors:** Joachim Seelig, Anna Seelig

## Abstract

The thermodynamics of pressure induced protein unfolding (denaturation) could so far not be directly compared with protein unfolding induced by temperature or chemical agents. Here we provide a new cooperative model for pressure induced protein denaturation, that allows the quantitative comparison of the three denaturing processes based on their free energy, enthalpy, entropy, and cooperativity. As model proteins we use apolipoprotein A-1 and lysozyme. The comparison shows that heat-induced unfolding is the most cooperative process. It is characterized by large positive enthalpies and entropies and (due to enthalpy-entropy compensation) a small negative free energies. Pressure denaturation is less cooperative. The entropies and enthalpies are less positive, and the resulting free energies are more negative. Chemically induced unfolding is least cooperative and shows the most negative free energies, in particular, if guanidinium hydrochloride (exhibiting a high binding affinity to certain proteins) is used as a denaturant. The three unfolding processes not only differ with respect to their cooperativity and the thermodynamic parameters, but also with respect to the volume changes. Whereas thermal and chemical denaturation increase the protein volume, denaturation by pressure reduces the protein volume, suggesting significant structural differences of the denatured proteins. Using cooperative models for protein analyses thus yields significant new insights into the protein unfolding/folding processes.

## Introduction

Proteins can be inactivated by heating,^1-5^ chemical agents,^6-8^ or high pressure.^9-10^ Inactivation by heating and chemical agents leads to unfolding and an incrase in volume, in contrast, application of high pressure leads to denaturation and a small reduction in protein volume.^11^ It can be expected that the three modes of inactivation lead to completely different unfolded structures. For the last 60 years, denaturation was analysed exclusively by a chemical equilibrium two-state model. This model assumes a chemical equilibrium between a native protein (N) and an unfolded protein (U). No intermediates are considered. The two-state model is proclaimed as the “most cooperative” unfolding model. However, the model contains no element of cooperativity and is, in fact, not different from simple cis - trans isomerisation. A cooperative model for protein denaturation was proposed by Zimm-Bragg^12^ in 1959, but was widely ignored. During the last 10 years, we have shown that the Zimm-Bragg theory can be successfully applied to protein denaturation.^13-17^ We have recently published comprehensive statistical-mechanical thermodynamic models for thermal^18-20^ and chemical^21-23^ unfolding. Here we extend our cooperative model to the analysis of protein denaturation under high pressure. We first compare the conventional chemical two-state model with a statistical two-state model. The focus of this work is however on a cooperative model of protein unfolding following, in principle, our previous proposals on thermal and chemical protein unfolding.^18-20, 23^

## Theory

The pressure dependence of protein stability is usually monitored by following a spectroscopic intensity I(p). The intensity change upon pressure application can be used to calculate an equilibrium constant K_NU_(p)^9^

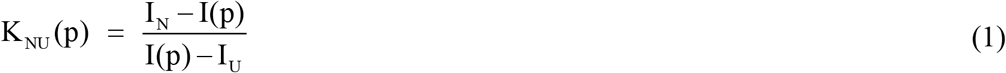

where I_N_ and I_U_ are spectroscopic intensities of the native and the unfolded (denatured) protein. With this definition, the native protein has a small equilibrium constant. The fraction of unfolded protein is given by

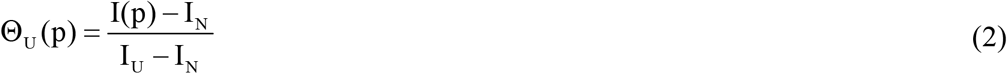

Three caveats should be noted. First, the spectroscopic intensity I(p) is only an indirect measure of protein unfolding, that is, it may correlate only approximately with changes in protein stability, protein function, or protein volume. Secondly, the intensity I_U_ may not represent the fully unfolded protein. The problem of incomplete protein unfolding under most experimental conditions has been discussed in reference.^24^ Thirdly, proteins are compressible even at low pressure. An example is Staphylococcal nuclease (SNase) (cf. Figure 5 in reference ^25^). The compressibility term **κ** = (1 / V)(∂V / ∂p) is not considered in the following. It is not reflected in the spectroscopic measurements (cf. Figure 4 in reference^26^).

### Chemical equilibrium two-state unfolding

In this model, pressure-unfolding is assumed to follow a two-state chemical equilibrium between a native N and an unfolded U protein. The pressure-dependence of the chemical equilibrium constant is given by^9^

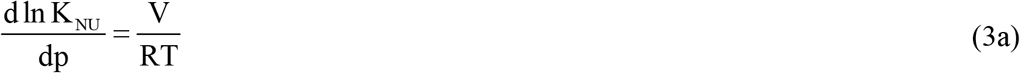

For the chemical equilibrium two-state model, a constant volume reduction ΔV_0_ is employed. Proteins are rather incompressible and ΔV_0_ is the decrease in protein volume upon applying pressure. Together with the pressure p_0_ at the midpoint of unfolding, ΔV_0_ is the second fit parameter to simulate the unfolding transition leading to

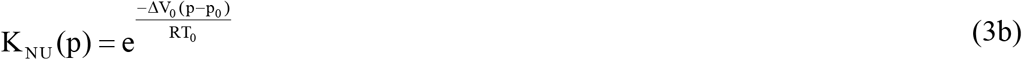

ΔV_0_ is negative and usually only 1% - 5% of the total protein volume.

The pressure dependence of the Gibbs free energy is

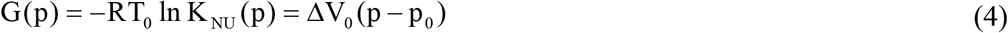

G(p) is a linear function of pressure. At p = p_0_ the equilibrium constant is K_NU_(p_0_) = 1. The native and the unfolded protein have the same concentration, and the free energy G(p_0_) = 0. For p < p_0_ and a negative ΔV_0_, the free energy becomes positive. The native protein at low pressure has a large positive free energy, which is against common expectations of a stable native protein.

The fraction of unfolded protein is

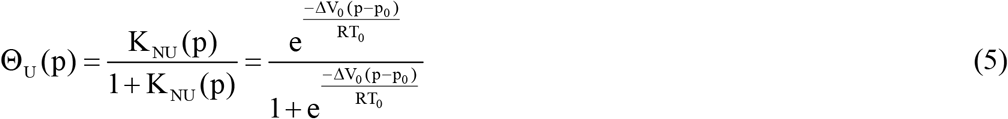

### Statistical-mechanical thermodynamic model of two-state unfolding

In this model, the protein flips between two energy states. The native protein is the reference state with energy E_N_ = 0. The unfolded state has the energy E_U_(p) = ΔV_0_(p−p_0_). The partition function of this two-state system is (for details see ^19-20, 22^)

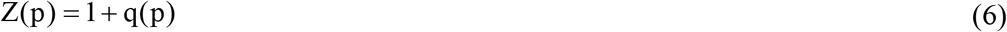

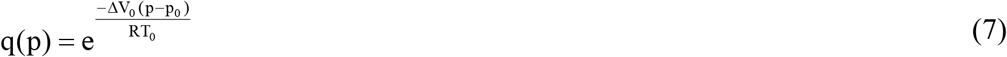

The fraction of unfolded protein is

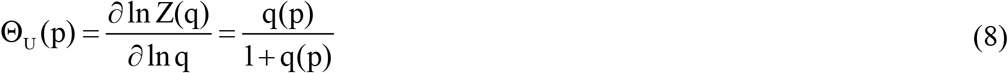

This equation is analogous to equation 5 of the chemical equilibrium model.^23^ However, there is a significant difference between the two models with respect to the pressure dependence of the free energy. The free energy of the statistical-mechanical model is

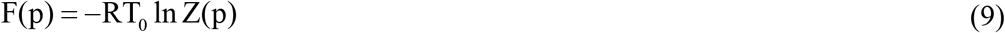

As long as the N-state is predominant, the free energy of the native state is close to zero. For p = p_0_, the free energy is not zero as in the chemical equilibrium model, but is F(p_0_) = −RT_0_ ln 2. For p > p_0_, the free energy F(p) coincides with G(p) of the chemical equilibrium two-state model.

### Cooperative multistate unfolding model

The cooperative multistate model assumes the interaction of ν molecular elements (e.g. amino acid residues or potein subdomains). Each element undergoes a transition from a native state “n” to an denatured state “u”. This could also be a small shift between subdomains reducing the total volume. The model has been applied previously to thermal and chemical protein unfolding.^18-20, 23^ In the present case of pressure unfolding the n → u transition is associated with a small volume change Δv_0_ per amino acid residue or subdomain. Δv_0_ is an average value over all cooperative elements. Each element is assumed to flip between a native state of energy 0 and an unfolded state of energy Δv_0_(p-p_0_). The cooperative interaction is characterised by a cooperativity parameter σ, which is also an average value. The partition function of the system is The fraction of unfolded protein is

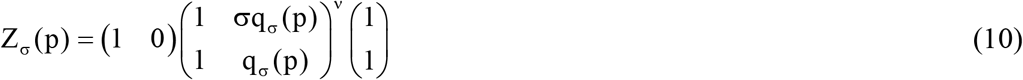

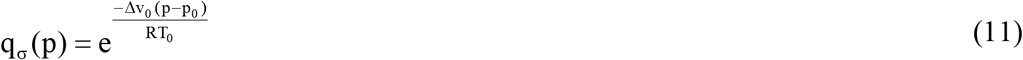

The fraction of unfolded protein is

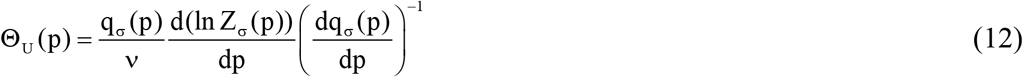

Finally, the pressure dependence of the free energy is

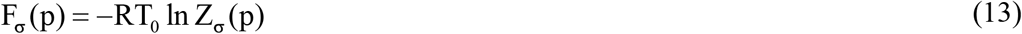

The fit parameters in the simulation of the thermodynamic properties are thus the cooperativity parameter σ, the volume change Δv_0_ per cooperative element and the number ν of cooperative elements participating in denaturation.

Figure 1 shows a comparison of the three unfolding models. The parameters are chosen to provide the best fit to lysozyme pressure-unfolding as described by figure 3 in reference.^27^ All three models provide a perfect fit to the experimentally observed sigmoidal unfolding transition Θ_u_(p). Moreover, the chemical equilibrium and the statistical-mechanical two-state model use the same set of parameters.

**Figure 1.**
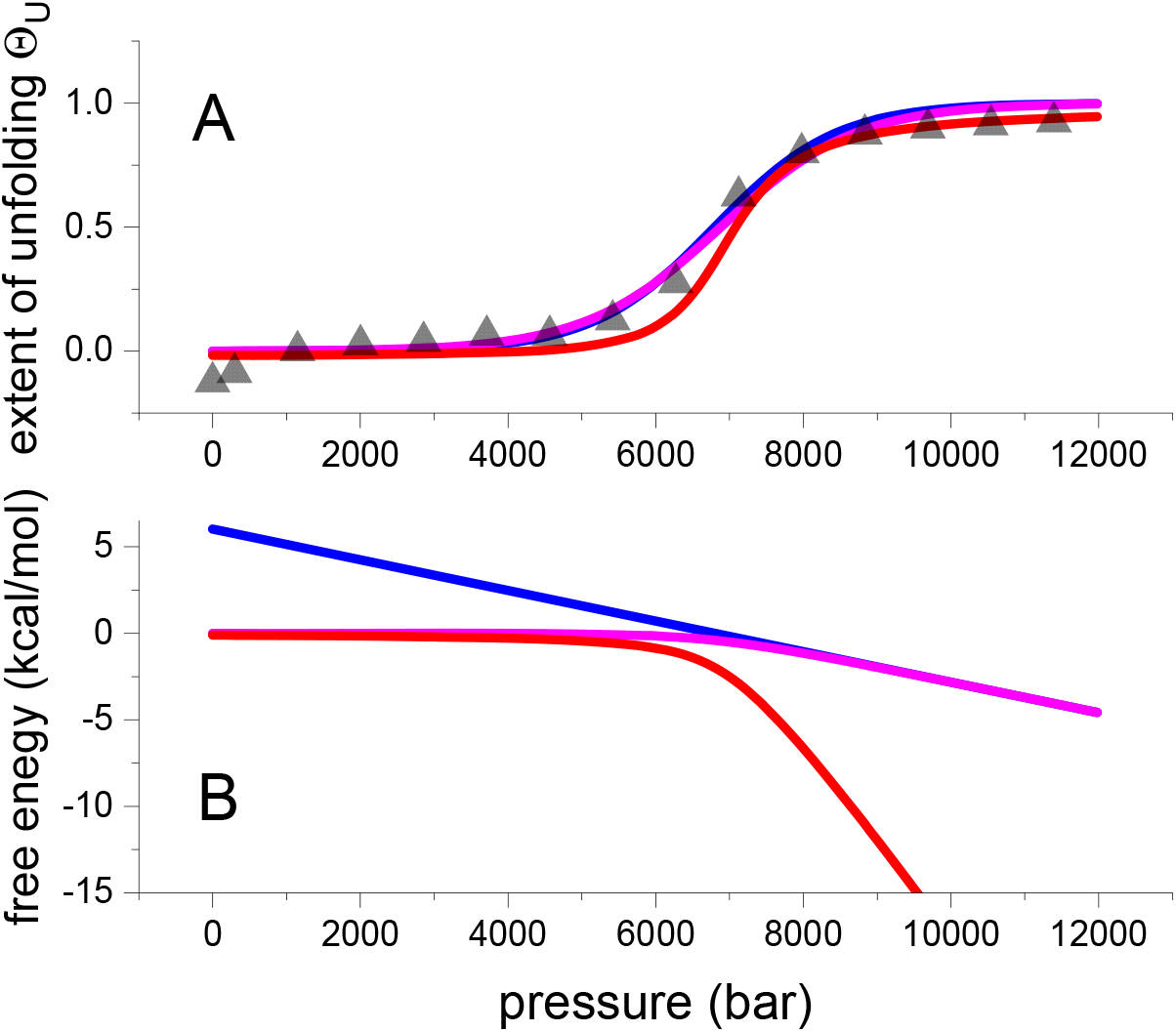
Comparison of three unfolding models. (A) (Δ) Experimental data of pressure-induced unfolding of lysozyme^27^ (details in figure 2). Extent of unfolding Θ_U_(p). The solid lines represent the best fit of each model to the experimental lysozyme data. Blue line: chemical two-state model with ΔV_0_= -37 mL/mol and p_0_ = 6800 bar. Magenta line: statistical-mechanical two-state model with ΔV_0_= -37 mL/mol and p_0_ =6800 bar. Red line: statistical-mechanical multistate cooperative model with volume change per amino acid residue Δv= - 2.0 mL/mol, ν = 129, p_0_ = 6800 bar. Cooperativity parameter σ = 5×10^−5^. (B) Free energy predictions of the three models. Blue line: chemical equilibrium two-state model (equation 4). Magenta line: statistical mechanical two-state model (equation 9). Red line: multistate cooperative model (equation 13). Same parameters as in panel A.

**Figure 2.**
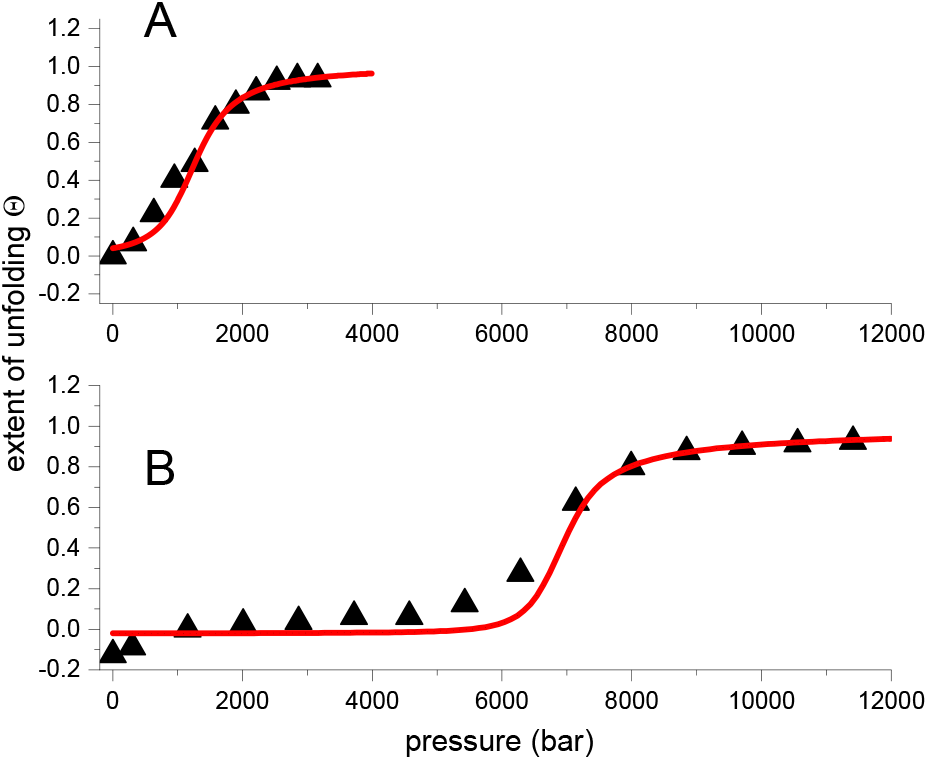
Pressure unfolding of ApoA1 and lysozyme at 25 °C. (A) Pressure unfolding of ApoA1 (black triangles). Experimental data obtained with FTIR-spectroscopy. Data taken from Mantulin and Pownall.^31^ Red line: multistate cooperative model (eq. 12). Cooperativity parameter σ = 1.6×10^−3^, volume per amino acid residue Δv = - 3.8 mL/mol. (B) Lysozyme. Experimental data (black triangles) measured with FTIR-spectroscopy, data taken from figure 3 of reference.^27^ Red line: multistate cooperative model (eq. 12). Cooperativity parameter σ = 5×10^−5^. Volume change per amino acid residue Δv = - 2.0 mL/mol.

**Figure 3.**
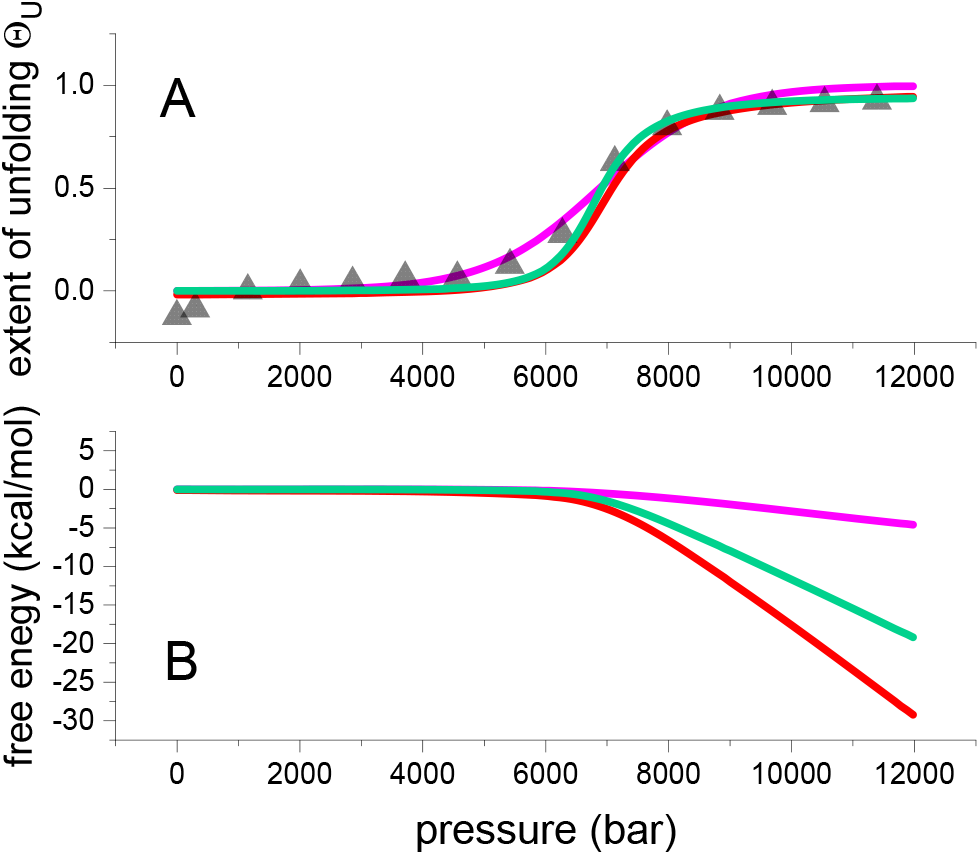
Pressure unfolding of lysozyme, modelled with 8 cooperative subdomains (green line), compared to the statistical-mechanical two-state model (magenta line) and the multistate cooperative model (with 129 aa) (red line). The parameters of the subdomain model are ν = 8 subdomains, ΔV = -20 mL/mol per subdomain and σ = 5×10^−2^.

However, as discussed, the pressure profiles of the free energies are quite different. The chemical equilibrium model predicts a linear dependence with a positive free energy of 6.0 kcal/mol at 1 bar (blue line in figure 1B). The statistical mechanical two-state model predicts a zero free energy for the native protein which becomes negative already at the midpoint of unfolding (F(p_0_) = −RT_0_ ln 2) (magenta line in figure 1B). At p > p_0_, the free energies of the two two-state models become identical. An again different result is obtained with the multistate cooperative model. The free energy of the native protein is also zero. However, the free energy of the denatured protein is distinctly more negative than the free energies of the other two models (red line in figure 1B). At p = p_0_, the free energy is - 2 kcal/mol at 25 °C. At 90% unfolding (p = 9100 bar) the predicted free energy of the multistate model is -12.5 kcal/mol.

### A multistate cooperative approximation

The above formalism can be approximated by a simpler expression, which avoids the matrix formalism. The partition function is approximated by the largest eigenvalue λ_0_ of the above matrix^28^. This leads to the following partition function

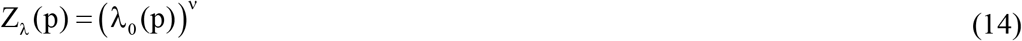

with

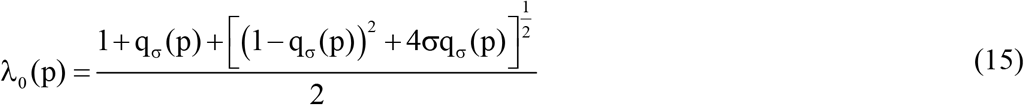

The fraction of unfolded protein is given by

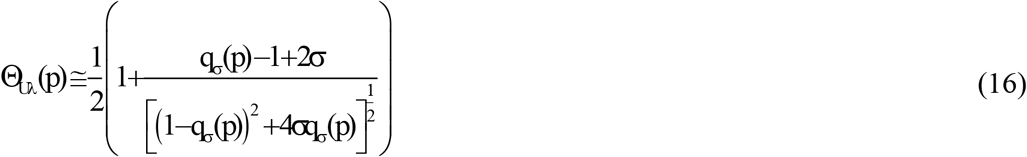

The free energy of unfolding is

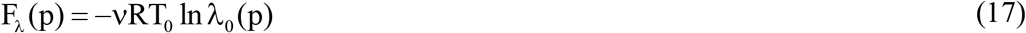

For pressures p ≫ p_0_, the following approximation is valid.

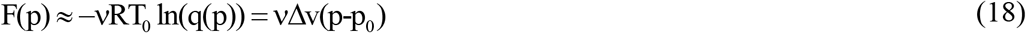

Good agreement between the approximation (equation 16) and the matrix approach (equation 12) was obtained in the analysis of experimental data obtained for lysozyme^27, 29^, SNase^26^ and ribonuclease (RNase) A.^26^

## Results

We apply the multistate cooperative model for thermal, chemical and pressure denaturation to the essentially linear α-helical apolipoprotein A-1 (ApoA1) and the compact globular lysozyme. Despite their different molecular weights (see below) the two proteins are comparable because the native-to-unfolded transitions involve 120 - 130 amino acid residues in both cases. The multistate cooperative model for high-pressure unfolding is closely related to analogous models for chemical^23^ and thermal^18^ unfolding.

### Apolipoprotein A-1

Human ApoA1 is an amphiphilic protein of 243 amino acid residues (28.16 kDa) that binds to lipid membranes and is an essential constituent in the formation of high density lipoprotein (HDL) particles that are required in the reverse cholesterol transport. The crystal structure of C-terminal truncated ApoA1 (183 amino acids) is almost 100% α - helical.^30^ ApoA1 in solution has an α-helix content of 50-60% (∼125 amino acid residues).^13-14^ The unfolding of ApoA1 is thus a cooperative sequential α-helix ⇄ random coil equilibrium, which can be described conveniently with the statistical-mechanical multistate theory.^13-14, 16^

### Lysozyme

Lysozyme (14.3 kDa) a 129-residue protein with ∼25% α-helix, ∼40% β-structure and ∼35% random coil in solution at room temperature.^15^ Upon thermal unfolding, the α-helix is almost completely lost and the random coil content increases to ∼60%.

The cooperative multistate theory is applied here to high pressure unfolding and the results will be compared to thermal^15^ and chemical^23^ unfolding, also described previously with the cooperative multistate theory.^18, 23^

### Pressure unfolding of apolipoprotein A-1 and lysozyme

Figure 2 compares the pressure unfolding of apolipoprotein A-1 (ApoA1) with that of lysozyme according to equation 12.

The relevant parameters are summarised in table 1.

Table 1

**Table 1.** Pressure denaturation of ApoA1 (experimental data from Matulin and Pownall^31^ and Lysozyme (experimental data from Smeller *et al*.^27^)

| *Parameters* | *Units* | *ApoA1* | *Lysozyme* |
| --- | --- | --- | --- |
| Number of amino acids, N<sub>aa</sub> | - | 123 | 129 |
| Midpoint pressure, p<sub>0</sub> | bar | 1150 | 6800 |
| Pressure at unfolding start, p<sub>ini</sub> | bar | 1 | 5000 |
| Pressure at 90% denaturation, p<sub>end</sub> | bar | 2501 | 9100 |
| Pressure diff. (p<sub>end</sub> - p<sub>ini</sub>) Δp | bar | 2500 | 4100 |
| Cooperativity parameter, σ | - | 1.6x10<sup>-3</sup> | 5x10<sup>-5</sup> |
| Volume change per aa residue, dv | mL/mol | -3.8 | -2.0 |
| Volume change per protein, ΔV<sub>0</sub> | mL/mol | -467 | -253 |
| Free energy change, ΔF | kcal/mol | -13.1 | -10.4 |
| ΔpΔV<sub>0</sub> | kcal/mol | 27.9 | 25.3 |
| Entropy change, ΔS | kcal/molK | 0.14 | 0.12 |

The open structure of ApoA1 with its linear α-helix is distinctly less stable than the more compact lysozyme. This is evidenced by the low midpoint pressure of 1150 bar. In contrast, the midpoint pressure of lysozyme is 6800 bar. The pressure range of unfolding is 2500 bar for ApoA1 but 4100 bar for lysozyme. Similar results for the pressure unfolding of lysozyme in H_2_O and D_2_O have been reported by Hedoux *et al*.^*29*^ Volume changes per amino acid residue are larger (−2.8 to -2.9 ml/mol) and the cooperativity parameters are σ =5 × 10^−5^ and σ =1 × 10^−4^, respectively.

Pressure unfolding of proteins is measured with spectroscopic methods that reflect volume changes only indirectly. The true volume reduction of the protein is not known. The multistate cooperative model offers a large range of different parameters to simulate the same experimental data (Δv, ν and σ). This is demonstrated in figure 3. The experimental data in figure 3A are identical to those in figures 1A and 2B and present the pressure unfolding of lysozyme. The green lines in figure 3 represents a subdomain unfolding model. It is assumed that lysozyme has 8 subdomains, which under pressure move closer together with a volume reduction of -20 mL/mol per subdomain. The cooperativity parameter is 5×10^−2^. Figure 3B demonstrates that the free energy of this subdomain model is between those of the two-state model and the multistate cooperative model with ν = 129 amino acid residues with a volume reduction of -3 mL/mol per amino acid residue.

As a general conclusion it thus follows that spectroscopic measurements of pressure unfolding are not specific enough to allow an unambiguous interpretation of the unfolding process. A direct measurement of the volume reduction would indeed be necessary.

### Heat denaturation of ApoA1 and lysozyme

Proteins are denatured by very low or very high temperatures. Depending on the protein, the denaturation temperature varies strongly. The method of choice to study thermal unfolding of proteins is differential scanning calorimetry (DSC).^32-33^ The heat capacity C_p_ is measured as a function of temperature and the important thermodynamic parameters of unfolding, that is, enthalpy H, entropy S and free energy G are obtained directly by integration of the experimental C_p_(T) values.^18-20^ Spectroscopic measurements yield in contrast only indirect information on thermodynamic properties and can deviate substantially from DSC measurements.^15^ Figure 4 displays DSC measurements of ApoA1 and lysozyme.

The simulation of the experimental data follows references.^18-20^ The simulation parameters are, h_0_, the enthalpy per amino acid residue, c_v_, the heat capacity per amino acid, and σ, the cooperativity parameter. The simulation parameters and other relevant thermodynamic results are summarised in table 2.

Table 2

**Table 2.** Thermal unfolding of ApoA1 (experimental data taken from Eckhardt *et al*.^16^) and Lysozyme (experimental data taken from Li-Blatter *et al*.^22^)

| *Parameters* | *Units* | *ApoA1* | *Lysozyme* |
| --- | --- | --- | --- |
| Number of amino acids, N<sub>aa</sub> | - | 123 | 129 |
| Midpoint temperature, T<sub>0</sub> | °C | 54 | 63 |
| Temp. at unfolding start, T<sub>ini</sub> | °C | 33 | 52 |
| Temp. at 90% unfolding, T<sub>end</sub> | °C | 77 | 77 |
| Temp. diff. (p<sub>end</sub> - p<sub>ini</sub>) ΔT | °C | 44 | 25 |
| Cooperativity parameter, σ | - | 5x10<sup>-5</sup> | 1x10<sup>-6</sup> |
| Enthalpy per aa, h<sub>0</sub> | cal/mol | 900 | 950 |
| Heat capacity per aa, c<sub>v</sub> | cal/molK | 8 | 7 |
| Molar heat capacity change, Δc<sub>p</sub> | kcal/molK | 2.72 | 2.48 |
| Conf. enthalpy change, ΔH<sub>conf</sub> | kcal/mol | 96 | 107 |
| Conf. entropy change ΔS<sub>conf</sub> | kcal/molK | 0.29 | 0.318 |
| Enthalpy change at 90% unfolding, ΔH | kcal/mol | 148 | 146 |
| Free energy change at 90% unfolding, ΔF | kcal/mol | -8.51 | -5.7 |
| Entropy change at 90% unfolding, ΔS | kcal/molK | 0.446 | 0.434 |

Thermal unfolding further confirms the lower stability of ApoA1 compared to lysozyme. The midpoint temperature of ApoA1 is 54 °C, that of lysozyme 63 °C. As the unfolding enthalpies of the two proteins are almost equal and T_0_ = ΔH_0_/ΔS_0_, the temperature difference is caused by differences in the unfolding entropy.

The unfolding temperature T_0_ of lysozyme depends on pH and increases with increasing pH.^33^ The present data (figure 4) refer to a PBS buffer at pH 2.5.^15^ Hedoux et al. studied lysozyme unfolding with DSC and Raman spectroscopy in water.^29, 34^ T_0_ is shifted to 74°C but the heat of unfolding is in broad agreement with the results given in table 2.

### Chemical unfolding of apolipoprotein A-1 and lysozyme

ApoA1 and lysozyme were denatured by adding increasing concentrations of guanidine HCl. Figure 5 shows the sigmoidal transition curves for chemical unfolding. Mantulin and Pownall measured the chemical denaturation of ApoA1 in parallel to pressure unfolding.^31^ Lysozyme chemical unfolding was reported by Ahmad and Bigelow.^35^

All data are summarised in table 3.

Table 3

**Table 3.** Chemical unfolding of ApoA1 (experimental data taken from Matulin *et al*.^31^) and Lysozyme (experimental data taken from Ahmad *et al*.^27^)

| *Parameters* | *Units* | *ApoA1* | *Lysozyme* |
| --- | --- | --- | --- |
| Number of amino acids, N<sub>aa</sub> | - | 123 | 129 |
| C<sub>gdm</sub> at midpoint of unfolding, c<sub>0</sub> | M | 1.07 | 4.1 |
| c<sub>Gdm</sub> at start of unfolding, c<sub>ini</sub> | M | 0.3 | 3.0 |
| c<sub>Gdm</sub> at 90% unfolding, c<sub>end</sub> | M | 1.6 | 5.0 |
| Conc. range of unfolding, Δc<sub>Gdm</sub> | M | 1.3 | 2.0 |
| Cooperativity parameter, σ | - | 2x10<sup>-2</sup> | 2x10<sup>-3</sup> |
| Binding constant of Gdm, K<sub>D</sub> | M<sup>-1</sup> | 0.95 | 0.245 |
| Free energy change, ΔF,<br>at 90% unfolding | kcal/mol | -30.9 | -14.1 |
| Entropy change, ΔS | kcal/molK | nd | 0.28 |
| Enthalpy change, ΔH | kcal/mol | nd | 70 |

The midpoint concentration for ApoA1 unfolding is only 1.07 M compared to 4.2 M for lysozyme. A low concentration of guanidine HCl is sufficient to denature ApoA1. It is the result of the better binding of guanidine HCl to ApoA1 (binding constant K_D_ = 0.9 M^-1^) than to lysozyme (K_D_ = 0.245 M^-1^). The cooperativity parameters are σ = 2×10^−2^ for ApoA1 and σ = 1×10^−3^ for lysozyme. The binding of the denaturant thus reduces the cooperativity in both proteins compared to pressure unfolding. Again, the unfolding of ApoA1 is less cooperative than that of lysozyme.

## Discussion

### Models for pressure unfolding of proteins

Pressure unfolding of proteins has usually been described by a chemical equilibrium two-state model. Here, we propose a multistate cooperative model as a flexible and physically more realistic alternative. In the limit of no cooperativity this model degenerates into a statistical-mechanical two-state model (figure 1, magenta line). Another limit of this model is a cooperative subdomain model (figure 3, green line). A small number of protein subdomains interact cooperatively and move closer together to reduce the protein volume.

The multistate cooperative model for protein unfolding follows the same principles as discussed recently for thermal^18-20^ and chemical unfolding.^23^ It is thus possible to compare pressure denaturation with thermal and chemical unfolding. Whereas heat and chemical unfolding lead to an increase in the protein volume, pressure denaturation leads to a decrease in protein volume. The three methods, pressure denaturation, heat unfolding and chemical unfolding thus likely lead to denatured proteins of different structure.

### Pressure unfolding versus heat unfolding and chemical unfolding

The common parameters of the different models are the cooperativity parameter б and the free energy involved in unfolding F(p) = - RT_0_ lnZ(p). These parameters are, on the one hand, characteristic for the nature of the proteins and on the other hand, yield information on the thermodynamics of differernt unfolding processes.^11^

The cooperativity parameter σ as a function of the free energy F(p) is illustrated in figure When comparing the three unfolding methods (figure 6), thermal unfolding is the most cooperative process (smallest σ), followed by decreasing cooperativity for pressure unfolding and chemical unfolding. ApoA1 displays a particularly low cooperativity in chemical unfolding. This can be explained by the strong binding of guanidine HCl to ApoA1. Guanidine HCl binds to most proteins with a binding constant of K_D_ ∼0.25 M^-1^.^23^ The binding constant to ApoA1 is much stronger, K_D_ ∼0.95 M^-1^. The strong binding opens the protein structure and reduces the cooperativity.

**Figure 4.**
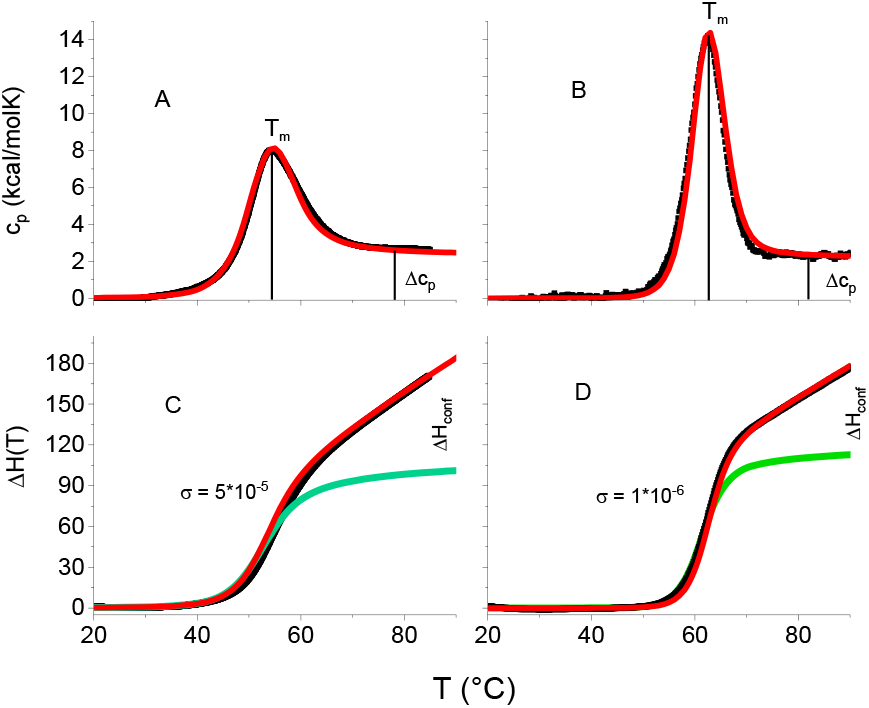
Heat denaturation of ApoA1 (A, C) and lysozyme (B, D) as measured by DSC. Black lines: about 2500 experimental DSC data points. Red lines: multistate cooperative model.^18-20^ Green lines: predicted enthalpy change ΔH_conf_ (T) of the conformational transition (A, B) Heat capacity C_p_(T). Midpoint temperatures T_0_=54 °C for ApoA1 (pH 7.4) and T_0_ = 63 °C for lysozyme (pH 2.5). (C, D). Unfolding enthalpies H(T) of ApoA1 and lysozyme obtained by integration of the heat capacities. The fit parameters and relevant thermodynamic parameters are summarised in table 2.

**Figure 5.**
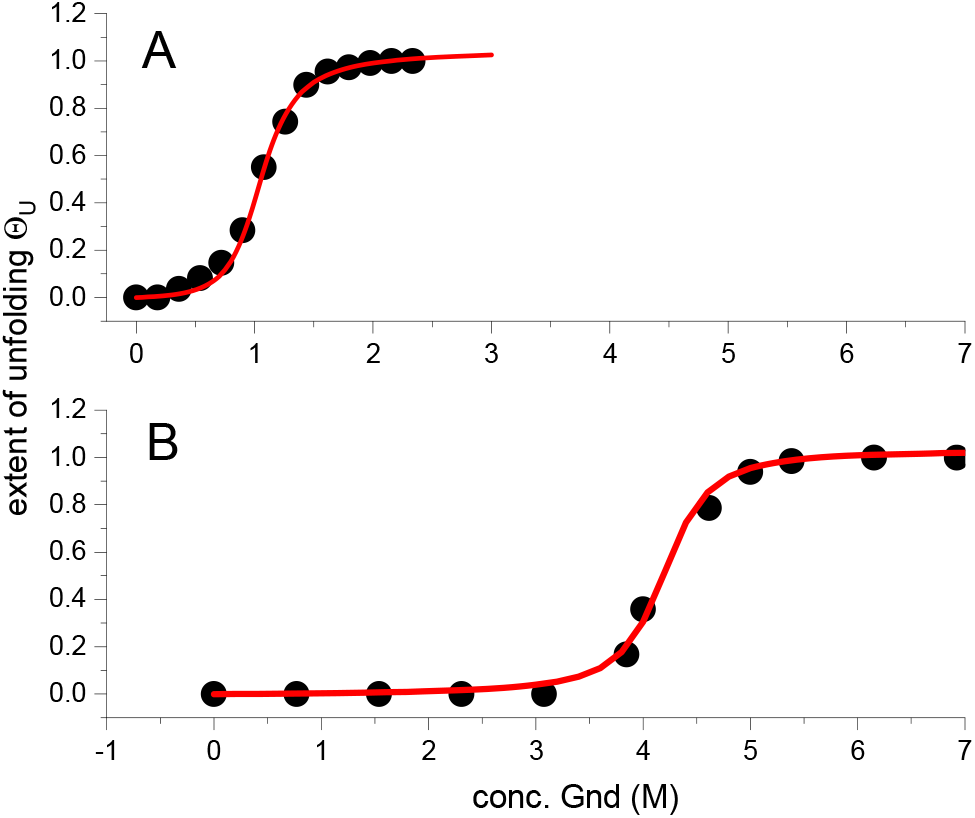
Chemical unfolding of ApoA1 and lysozyme. (A) ApoA1 (•). Experimental data^31^. Midpoint concentration c_0_ = 1.07 M. Cooperativity parameter σ = 2×10^−2^. Guanidine HCl binding constant K_D_ = 0.95 M^-1^. (B) Lysozyme. (•) Experimental data.^35^ Midpoint concentration c_0_ = 4.2 M. Cooperativity parameter σ = 2×10^−3^. Guanidine HCl binding constant K_D_ = 0.245 M^-1^.

**Figure 6.**
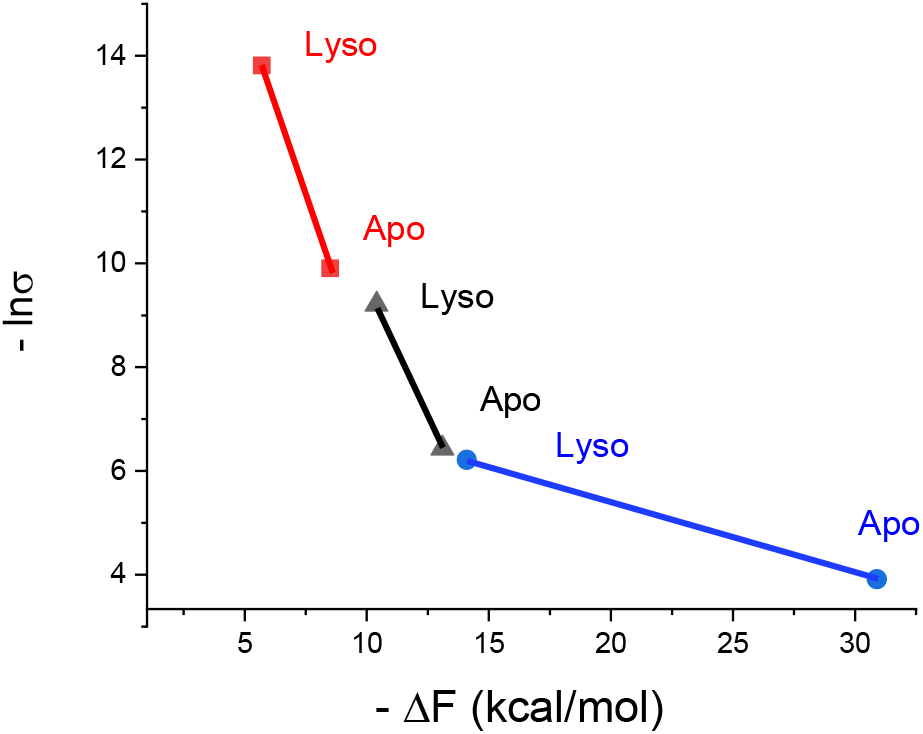
Comparison of the three unfolding processes using common parameters. The cooperativity, expressed as –lnб, is plotted as a function of the free energy released upon unfolding for ApoA1 (Apo) and lysozyme (Lyso). Temperature-induced unfolding (red), pressure-induced unfolding (black) chemical unfolding (blue).

Heat unfolding yields the smallest negative free energy change, followed by pressure denaturation, and chemical unfolding. An approximately linear relationship between the free energy change and the free energy of nucleation −RT_0_ ln **σ** (see Figure 6) is observed for thermal and pressure unfolding. Lysozyme is generally slightly more cooperative and yields a less negative free energy change than ApoA1.

Interesting molecular insight is provided by considering the entropy change. The largest entropy change is observed for thermal unfolding with ΔS ∼ 0.44 kcal/mol for both proteins (table 1). Chemical unfolding of lysozyme results in a lower entropy change of ΔS ∼ 0.28 kcal/mol,^23^ (table 3) because part of the entropy gained upon unfolding is lost again upon binding of denaturants. The smallest entropy change is produced by pressure unfolding with ΔS ∼ 0.14 kcal/mol for ApoA1 and ∼ 0.09 kcal/mol for lysozyme (table 2). This suggests that the structural changes induced by high-pressure are much smaller than those obtained with thermal and chemical unfolding.

### Volume changes upon pressure application

Denaturation of proteins by heat or chemical agents, produces an increase in protein dimensions. This can be measured directly, for example, by dynamic light scattering.^11^ In contrast, pressure, denaturation of proteins, leads to a small decrease in protein volume. Unfortunately, this volume reduction has not yet been measured directly, but follows from the interpretation of spectroscopic unfolding measurements with unfolding models. For lysozyme the two-state models predict a volume reduction of -37 mL/mol, the subdomain model 8 x -20 = -160 mL/mol, and the multistate cooperative model 129 x -2 = - 258 mL/mol. All three models describe the sigmoidal unfolding transition perfectly well.

We have analysed pressure isotherms of ApoA1^31^, lysozyme^27, 29^, SNase^26^ and RNase^26^ with the two-state models and the multistate cooperative model. The average volume change of 6 pressure isotherms of these proteins in the two-state analysis was -60 ± 16 mL/mol. For the multistate cooperative model, the average volume change per amino acid residue was -2.9 ± 0.7 mL/mol, which results in -350 ± 80 mL/mol as the average volume change of the whole proteins.

As there are no direct volume measurements, we estimate volume changes from the experimentally available protein compressibility. The protein compressibility 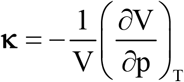 was measured as κ = 2×10^−11^ - 8×10^−11^ Pa^-1^ for SNase.^25^ The predicted volume change is thus ΔV(p) = −V_0_ (1− e^−**κ**p^). For a volume of V_0_ ∼ 10 L/mol (e.g., SNase) at a pressure of 3000 bar the predicted volume change, calculated with the compressibility κ = 8×10^−11^ Pa^-1^ is ΔV = -259 mL/mol. This calculation, while only approximate, shows that large volume changes can be expected and are consistent with the multistate cooperative model.

A molecular interpretation of volume changes under pressure has been given.^36^ Pressure reduces the void volume, but enhances the hydration sphere of the protein (see figure 8 in reference^36^). For a protein with 100 - 200 amino acid residues a volume change of∼ -100 mL/mol is predicted.^36^

The different protein behaviours under high temperature and high pressure result to some extent to changes from the dielectric constant, ε, of water. An increase in temperature reduces the dielectric constant of water, whereas in increase in pressure enhances the dielectric constant of water. Figure 7 shows the cooperativity of unfolding as a function of the dielectric constant of water at the midpoint of temperature and pressure unfolding. Increasing the temperatue reduces the polarity of water, which facilitates water penetration into the more hydrophobic protein interior. An increase in pressure enhances the polarity of water. This reinforces hydrogen bonding between water molecules and reduces their penetration into the protein interior, leading to more pronounced hydration shells of the protein.

**Figure 7.**
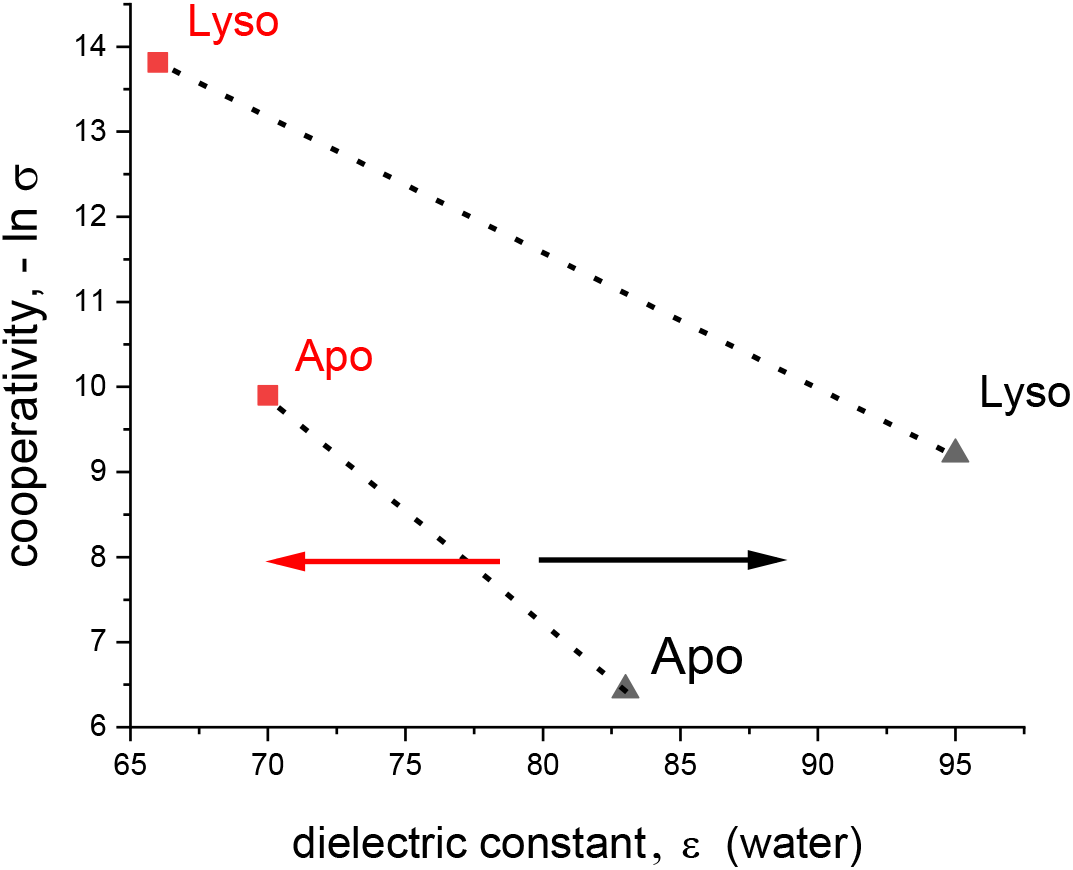
The cooperativity of unfolding (– ln σ as a function of the dielectric constant, ε of water at the specific temperatures T_0_ (red) and pressures p_0_ (black) for ApoA1 and lysozyme. At room temperature and atmospheric pressure the dielectric constant of water is ε ∼ 80. Increasing the temperature reduces the dielectric constant of water (red arrow). Increasing the pressure enhances the dielectric constant of water (black arrow).

## Conclusion

Protein folding/unfolding is a cooperative process. We propose a truly cooperative model for the high pressure unfolding of proteins. The model may involve the cooperative interaction of all amino acids of a particular protein, but can also handle the cooperative volume reduction of a small number of subdomains. In the limit of no cooperativity the model degenerates into a statistical-mechanical two-state model. The latter model is quite similar to the well-known chemical equilibrium two state model, but predicts a different temperature profile of the free energy. In the multistate cooperative model the native protein is the reference state with free energy zero.

The new model is built on the same principles as described for the cooperative unfolding induced by heat or chemical agents. It is thus possible for the first time to draw a comparison between the three methods of protein denaturation. Thermal unfolding is the most cooperative process, followed by pressure unfolding and chemical unfolding. On the other hand, the change in free energy is less negative for thermal unfolding than for pressure unfolding and chemical unfolding. Particularly interesting is a comparison of the entropies. Pressure unfolding displays the smallest enthalpy change, which is probably indicative of the much smaller structural changes induced by pressure compared to heat and chemical agents. Examples are given for an essentially helical protein (ApoA1) and a globular protein (lysozyme). Although, the differences between these two proteins are small, the data provide insight into the specific unfolding processes.

